# Personalized *in-silico* drug response prediction based on the genetic landscape of muscle-invasive bladder cancer

**DOI:** 10.1101/2020.05.22.101428

**Authors:** Friedemann Krentel, Franziska Singer, María Lourdes Rosano-Gonzalez, Ewan A. Gibb, Yang Liu, Elai Davicioni, Nicola Keller, Daniel Stekhoven, Marianna Kruithof-de Julio, Roland Seiler

## Abstract

In bladder cancer (BLCA) there are, to date, no reliable diagnostics available to predict the potential benefit of a therapeutic approach. The extraordinarily high molecular heterogeneity of BLCA might explain its wide range of therapy responses to empiric treatments. To better stratify patients for treatment response, we present a highly automated workflow for *in-silico* drug response prediction based on a tumor’s individual multi-omic profile. Within the TCGA-BLCA cohort, the algorithm identified a panel of 21 genes and 72 drugs, that suggested personalized treatment for 94,7% of patients - including five genes not yet reported as biomarkers for clinical testing in BLCA. The automated predictions were complemented by manually curated data, thus allowing for accurate sensitivity- or resistance-directed drug response predictions. Manual curation revealed pitfalls of current, and potential of future drug-gene interaction databases. Functional testing in patient derived models and/or clinical trials are next steps to validate our *in-silico* drug predictions.

## Introduction

More than three decades of development in empiric systemic therapies have passed without remarkably changing the numbers of deaths caused by bladder cancer (BLCA).^1^ As of 2017, BLCA is the seventh most common cancer and annually affects more than 450.000 people worldwide, resulting in almost 200.000 deaths.^2^ Recent advancements include immunotherapies in BLCA, which have brought the first significant progress in survival since platinum-based chemotherapy regimens were introduced in the mid 1980s.^3^ However, in recent years the five-year overall survival has only been improved by 5% - and the effects of immunotherapies are still to be evaluated.^4^

The molecular diversity of BLCA might be one of the major reasons for the low response rates of BLCA to systemic agents. BLCA is among the cancer types with the highest total mutational burden.^5^ Moreover, about two thirds of BLCA are exposed to APOBEC mutagenesis, which increases the mutational diversity within each individual cancer during development. At the transcriptomic level, BLCA can be divided into molecular subtypes and clinical implications of these sub-classifications have already been suggested.^6^ Nevertheless, transcriptome-based subclassifications alone do not seem to be optimal for selecting the right therapeutic approach, as subtypes are still highly heterogeneous and do not incorporate the entire genomic landscape of BLCA tumors.

To facilitate a biomarker driven personalized therapy approach in BLCA, the first step is the identification of clinically relevant and actionable genomic alterations. Thus, in this study we used the TCGA-BLCA cohort as a starting point and aimed to determine potentially actionable genes and associated drugs based on the mutational landscape of muscle invasive bladder cancer. Moreover, we integrated existing evidence from other cancer entities and transcriptomic data to enrich the genomic-based information. With this study we succeeded in broadening the horizon of agents to drive pretreatment diagnostics and decision making from an empiric towards an individualized marker driven therapy strategy.

## Results

### Variant calling, quality control and variant filtering

We first performed variant calling from whole-exome sequencing (WES) data from the TCGA-BLCA cohort (Cell, 2017). The derived single nucleotide variants (SNVs) and copy number variations (CNVs) served as a base for a query in “The Drug Gene Interaction Database” (DGIdb)^7^ to identify variants with non-curated* DGI information (* = all DGIs not yet curated by manual curation or not derived from an expert curated DGI database will hereafter be called “non-curated” to highlight their level of evidence). Intending to target clinically relevant mutations only, we performed a prediction on the variant effect on the protein function and included only those variants with potentially damaging effect. Furthermore, only variants with a high impact, e.g. missense or stop-gain variants, were retained, thereby removing variants with likely limited effect on the protein. Subsequently, we filtered by the cohort prevalence of affected genes and the type of drug-trials the DGIs had been derived from. Finally, selected DGIs not present in an expert curated database were manually curated (see **Fig. 1** and **Methods** for a detailed description) and transcriptome based drug-response scores (DRS) were calculated, if available.

**Fig. 1.**
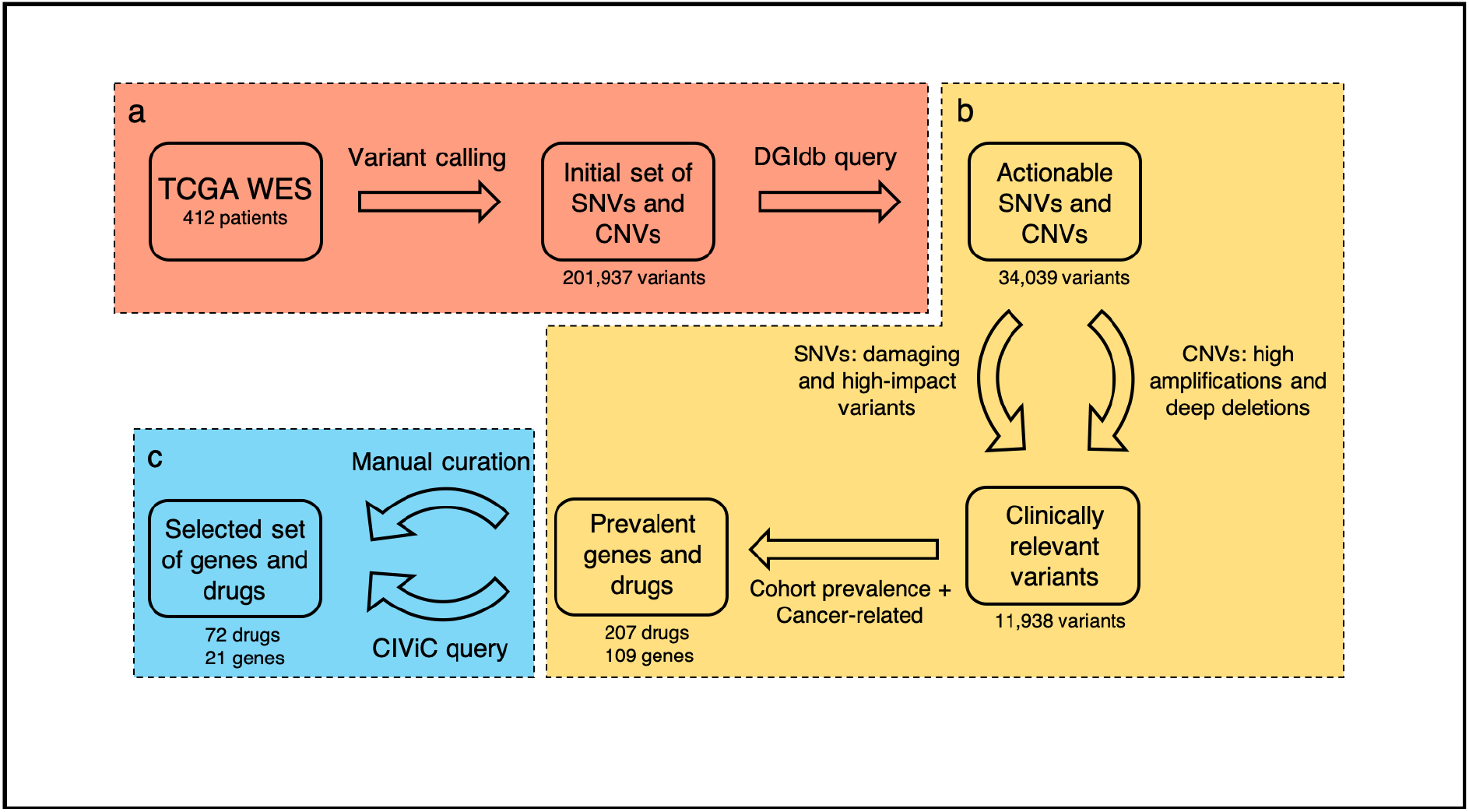
Workflow design - the diagram depicts the key steps of our *in-silico* drug prediction. **a** After variant calling in whole-exome sequencing (WES) data from the TCGA bladder cancer cohort, a query was performed on the “Drug Gene Interaction database” (DGIdb) to identify targetable genomic alterations amongst the discovered mutations. **b** During the subsequent fully automated filtering phase, further selection criteria were applied. **c** In the semi-automated filtering phase manual curation complemented the automated query of the expert curated CIViC database, resulting in the final set of genes and drugs.

### Variant calling compared to previous TCGA analysis

Variant calling from the 412 patients’ tumor genomes resulted in an unfiltered variant count (UVC) of 201,937 genomic variations. Quality metrics identified a high overlap with previous variant calling from the TCGA-BLCA cohort e.g. with Robertson et al. (Cell, 2017) With a median agreement of 98.3%, the identified SNVs in particular were highly similar to the previously documented analyses, despite using a more conservative approach than Robertson et al. (Cell, 2017). For identified CNVs the median agreement with previously reported CNVs was 84.6%. The lower agreement compared to SNV calls can be explained for example by the larger difference in the workflows used for CNV calling compared to SNV calling.

### Variant and drug filtering

DGIdb is the largest cumulative database for DGIs. It works as a widely used open source tool to collect drug-gene interactions by text mining from public resources, such as publications and databases,^7^ of which several, but not all, are expert-curated. By querying DGIdb, we identified 34,039 SNVs and CNVs with existing references for non-curated DGIs (in total 16,9% of UVC).

Before DGIdb filtering, tumor samples showed a wide range of mutational load from 1 to 7320 SNVs (median 293). The vast majority of samples presented drug-related SNVs after filtering, ranging from 0 to 574, with a median of 23 mutations. (see **Fig. 2a**). The subsequent mutational impact prediction for the SNVs identified a large overlap of the high-impact damaging variants with the total amount of all non-synonymous SNVs (see **Fig. 2b**). We observed a minor decrease in CNV count by filtering for deep deletions and amplifications, respectively. The CNV count per sample changed from 0 to 595 (median 52) prior to filtering, to 0 to 243 (median 44) after (see **Fig. 2c**). Copy number gains and amplifications outweighed copy number losses and deep deletions (see **Fig. 2d**).

**Fig. 2.**
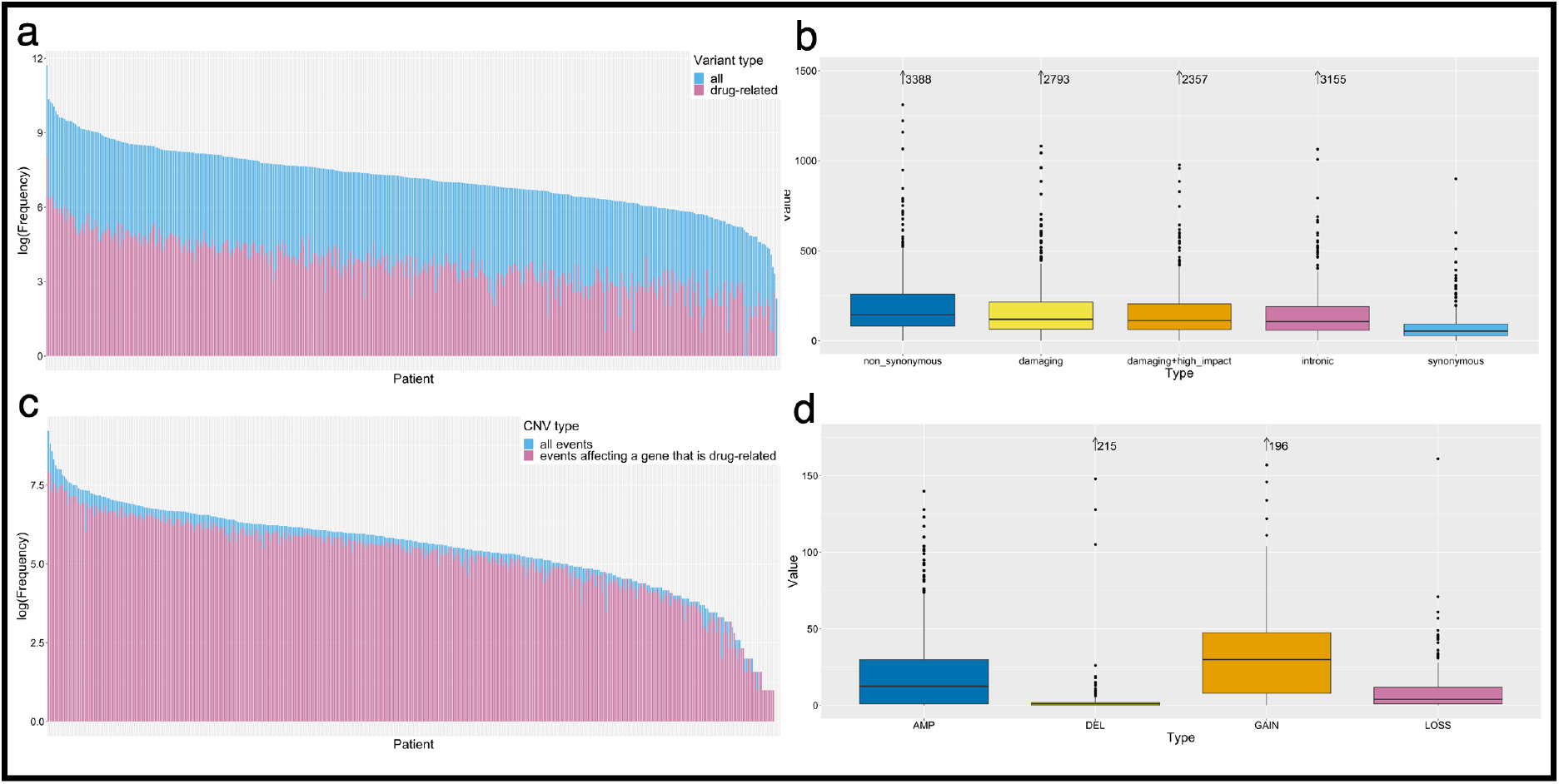
Filtering shows distinct effects on the SNV/CNV selection. **a** Barplot illustrates the log-scaled mutation load per sample for non-synonymous mutations, before (red bars) and after (blue bars) filtering for provided drug-gene interactions. Bars are overlying, not stacked. **b** Boxplots depict the distribution of types of SNVs before filtering. **c** The barplot shows the differences of CNV load per sample throughout filtering. The overlying bars represent CNV load before (red bars) and after (blue bars) filtering for non-curated drug-gene interactions. **d** Boxplots illustrate the type of CNV (DEL = deletion, 0 copies; LOSS = copy number loss, 1 copy; GAIN = copy number gain, 3-4 copies; AMP = amplification, >4 copies).

Next, we assigned the non-curated DGIs to the individual samples. This resulted in a “drug load” per sample, representing all predicted drugs with a gene dependent recommendation. **Fig. 3** illustrates the effect of each step of filtering on the “drug load”. The combined “drug load” considering all drugs for all SNVs and CNVs varied from 0 to 1896 (median 1422).

**Fig. 3.**
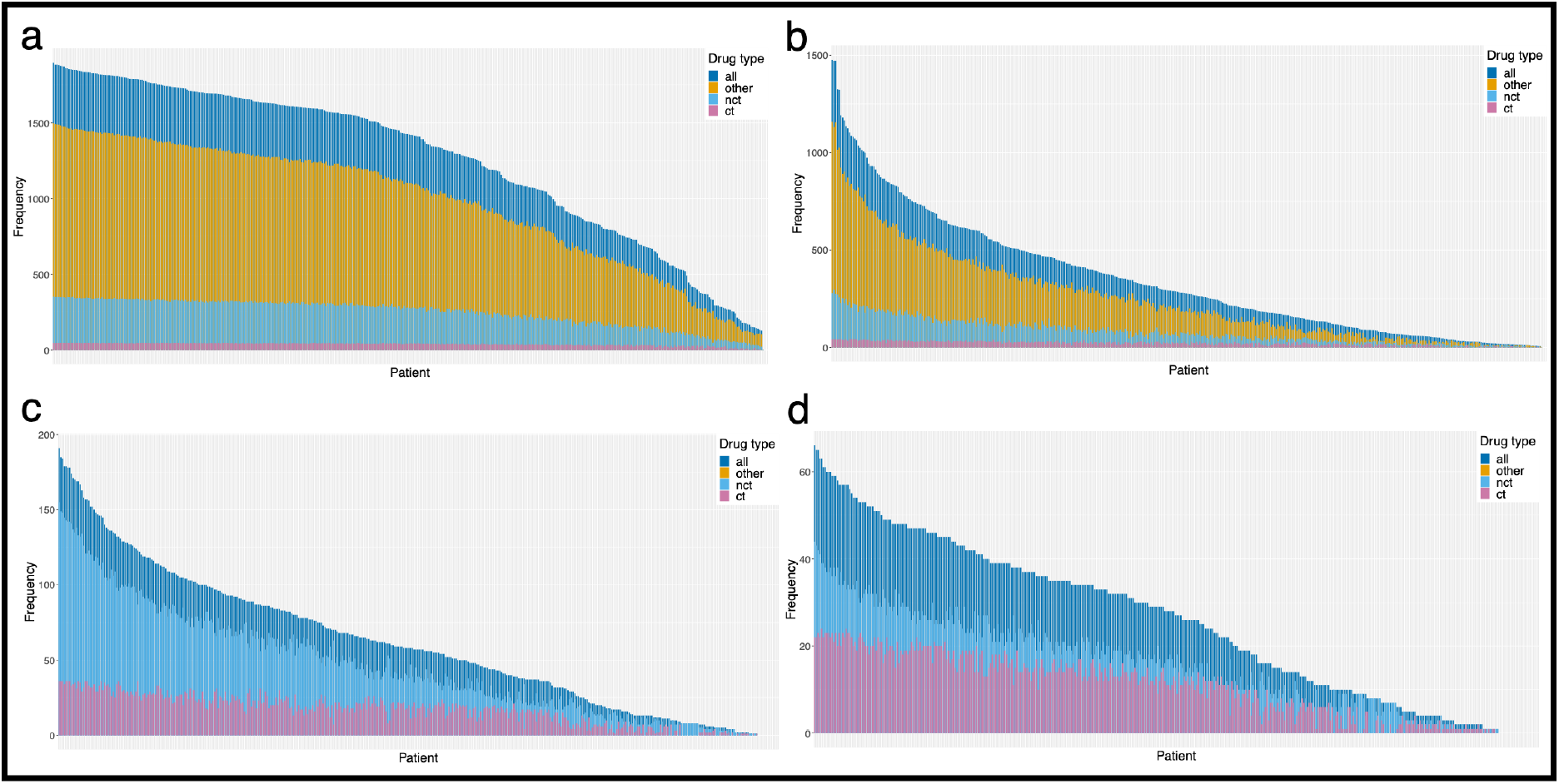
Development of the “drug load” along the filtering process. Each bar in **a-d** depicts a “drug load” per patient. Bars are not stacked, but overlying. Note that the y-axes change throughout the filtering. Drugs tested in bladder cancer trials or trials generally including solid tumors (ct) were distinguished from drugs used in general cancer trials (nct) and from medications not related to cancer in general. In **a-d** purple bars illustrate the total drug load per sample. Drugs with no reference to cancer therapy are illustrated in yellow (other). **a** Barplot illustrates the “drug load” for all SNVs and CNVs after querying DGIdg before any filters were applied. In **b** “drug load” after filtering for clinically relevant variants is depicted. In **c** cohort prevalence filters for variants were applied and drugs with no reference to cancer therapy were excluded. **d** Barplot illustrates the final “drug load” for curated drug-gene interactions after integration of CIViC and manual curation results.

Filtering DGIs for the selected variants with predicted clinical relevance (high transcriptome impact score) reduced the “drug load” to 0 to 1478 (median 274) drugs per sample. Applying cohort prevalence filters and excluding drugs without relation to cancer therapy resulted in 0 to 191 (median 57) drugs per sample. Manual curation and the CIViC query created a final total “drug load” of 0 to 66 drugs (median 27) per sample.

From the remaining 11,938 variants present across the cohort (5,9 % of UVC), for further investigation we demanded a variant cohort prevalence of 15%, which resulted in a set of 109 genes and 207 drugs. Thus, of the initial 4117 DGIs 91,2% were dismissed during automated filtering before manual curation and CIViC query. A majority of drop out resulted from both variant and drug related criteria. Drugs not being related to cancer therapy caused the dismissal of 17,5% of DGIs. Low cohort prevalence and non-damaging variants resulted in the removal of a further 18,3% (see **Fig. 3 and 4a**).

**Fig. 4.**
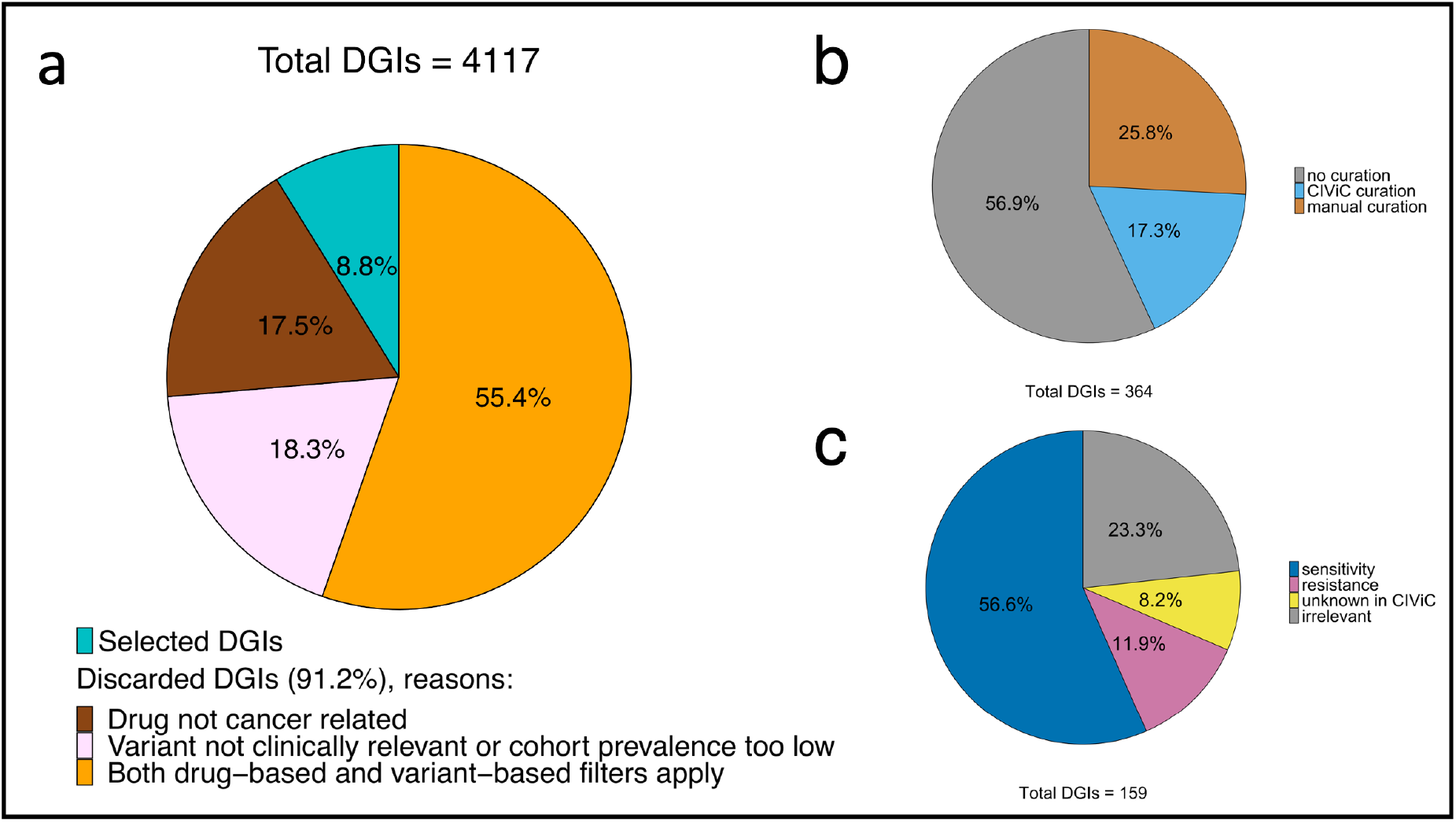
Influence of automated and manual filtering on the number of drug-gene interactions (DGIs). **a** The pie chart illustrates the fraction of DGIs filtered in an automated fashion, grouped by filter criteria. In **b**, the fraction of DGIs that underwent curation or CIViC query is shown (note that not all available DGIs were manually curated, as this was beyond the scope of this study), whereas in **c**, the respective assessment categories of the remaining DGIs with manual curation and CIViC support are depicted. Note that combination treatments are excluded in this plot, since they all refer to the same DGIs and would thus be counted twice.

### Automation / Manual Curation / CIViC Integration

Up to this point in our investigation the workflow was fully automated (see **Fig. 1 a and b)**. The selected DGIs resulted in a set of 109 genes and 207 associated drugs. However, the level of evidence of selected DGIs can be separated into two groups. The first group was supported by CIViC (“Clinical Interpretation of Variants in Cancer”; https://civicdb.org) DGI references. CIViC is an open source database for expert curated DGI references, which we integrated into our workflow to be automatically queried.^8^ The second group was compiled of non-curated DGIdb entries and lacks information on the actual support of the treatment. To base our *in-silico* predictions on proven DGIs only, we conducted a manual curation of non-curated DGIs (see **Methods - Manual Curation**). Note that we selected all SNV dependent non-curated DGIs for manual curation. Manual curation is very time consuming and we expected similar methodological results for CNV dependent DGIs. The non-curated SNV dependent DGIs selected for manual curation represent 25,8% of all selected DGIs (see **Fig. 4b)**. It can be expected that the fraction of non-curated DGIs will decrease over time, with more comprehensive information available in databases such as CIViC.

Interestingly, manual curation uncovered 62% of all investigated DGI references to be irrelevant, due to missing evidence of single agent efficacy or missing references to drug or gene itself. Commonalities of sources without proper reference to the DGI of interest include administration of the drug as part of a therapy regimen (which does not allow for evaluation of the single agent efficacy) and exclusive citations of either drug or gene in the list of references. In respect to all manually curated DGIs, 23.3% proved to be irrelevant due to exclusion in manual curation.

By extracting information concerning the direction of the DGI (sensitivity/resistance) from the DGI references, we were able to create binary predictions of response direction. While a majority of 55.6% of DGIs described sensitivity reactions, 11.9% of DGIs referred to resistance and 8.2% contained conflicting evidence e.g. in case of variant dependent contrary response prediction within one gene (see **Fig. 4c**). Additionally, we used information from the mined references to annotate DGIs and discovered 10 reasonable combinational therapies. Finally, filtering out DGIs neither supported by CIViC information nor by manual curation, reduced the number to 21 final genes and 72 drug/drug combinations, respectively.

### Integration of transcriptomic data

To base our predictions on a second data level we used *in-vitro* drug sensitivities and protein expression data from the NCI-60 cell lines to create drug response scores (DRS). The transcriptome dependent DRS were applied on each individual sample. DRS were measured for 26% (19/72) of the finally selected drugs. Note that a positive DRS indicates that the sample responded favorably to the tested drug, i.e. a positive DRS is interpreted as sensitivity. However, it is important to note that a negative DRS does not implicitly mean that the cell line was resistant to the applied drug, rather that no sensitivity could be observed. Thus, a negative DRS is interpreted as “no evidence of sensitivity”. The transcriptomic data based DRS provided a second level of information to weigh the inferred *in-silico* drug predictions. Additional evidence on the transcriptome level is an even stronger indication that (i) the variant observed on the genomic level has an impact and (ii) that the drug actually has the predicted effect. Thus, a DGI with directional support (sensitivity/resistance) on the genomic (SNV and/or CNV - derived) level, as well as support on the transcriptomic level, has a higher reliability compared to a DGI with support of only one level. This is illustrated in **Fig. 5**, where DGIs per sample are categorized based on their level of evidence for drugs with DRS support.

**Fig. 5.**
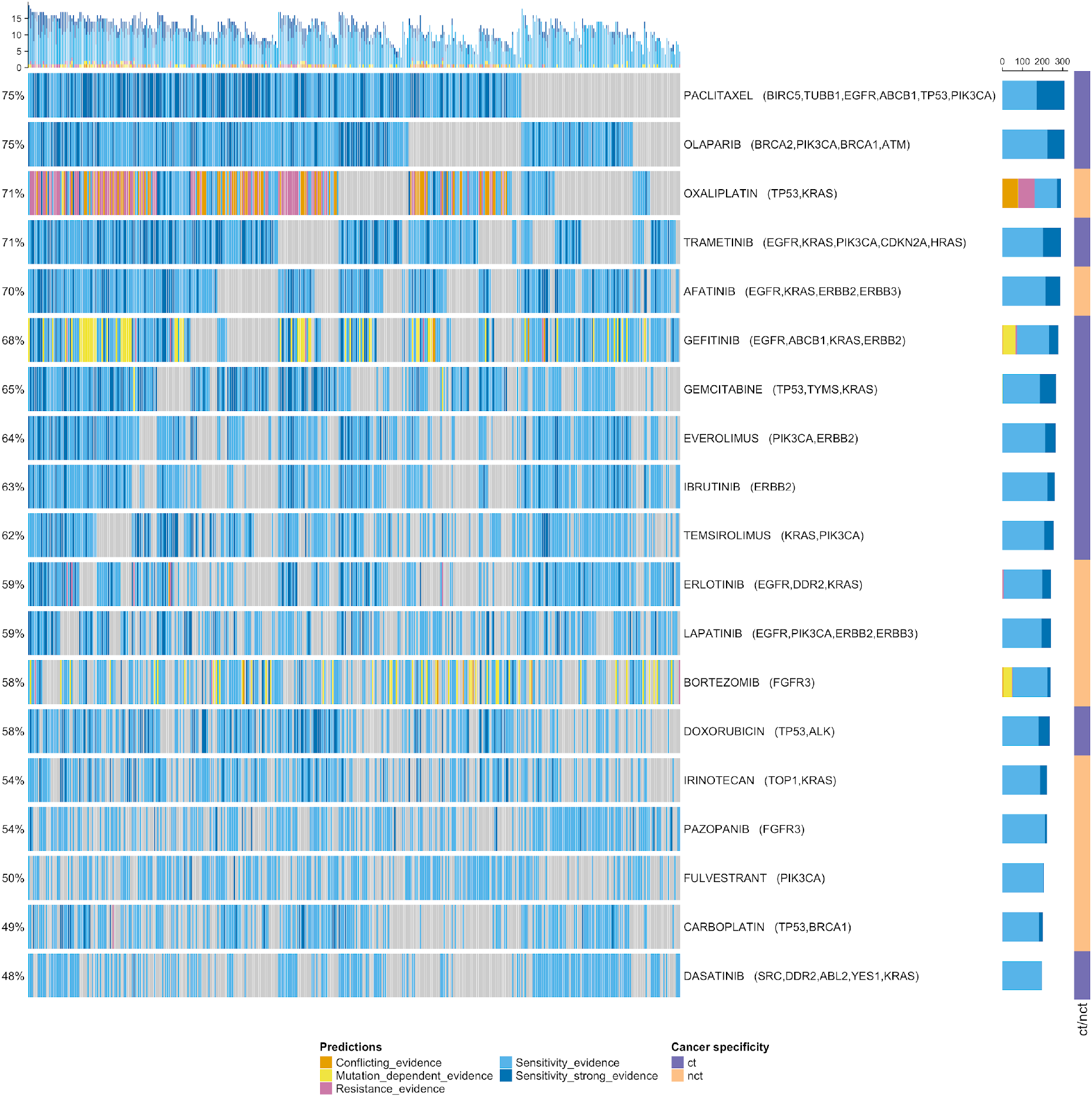
Oncoprint showing weighted evidence across samples based on DRS and curation support. Conflicting_evidence: resistance prediction + DRS positive; Mutation_dependent_evidence: gene_dependent prediction + DRS positive / gene_dependent prediction + no DRS; Resistance_evidence: resistance prediction + no DRS; Sensitivity_evidence: sensitivity_unspecific prediction + no DRS / sensitivity prediction + no DRS / no_genomic + DRS positive; Sensitivity_strong_evidence: sensitivity_unspecific prediction + DRS positive / sensitivity prediction + DRS positive.

### Final drug and gene panel

Based on the previously described steps, we identified a set of 21 actionable genes and 72 corresponding drugs or drug combinations as promising target-driven therapy suggestions for muscle invasive bladder cancer (see **Fig. 6**). In total 94.7% of TCGA-BLCA samples could be covered by our set of selected drugs and genes by being represented with at least one predicted DGI.

**Fig. 6.**
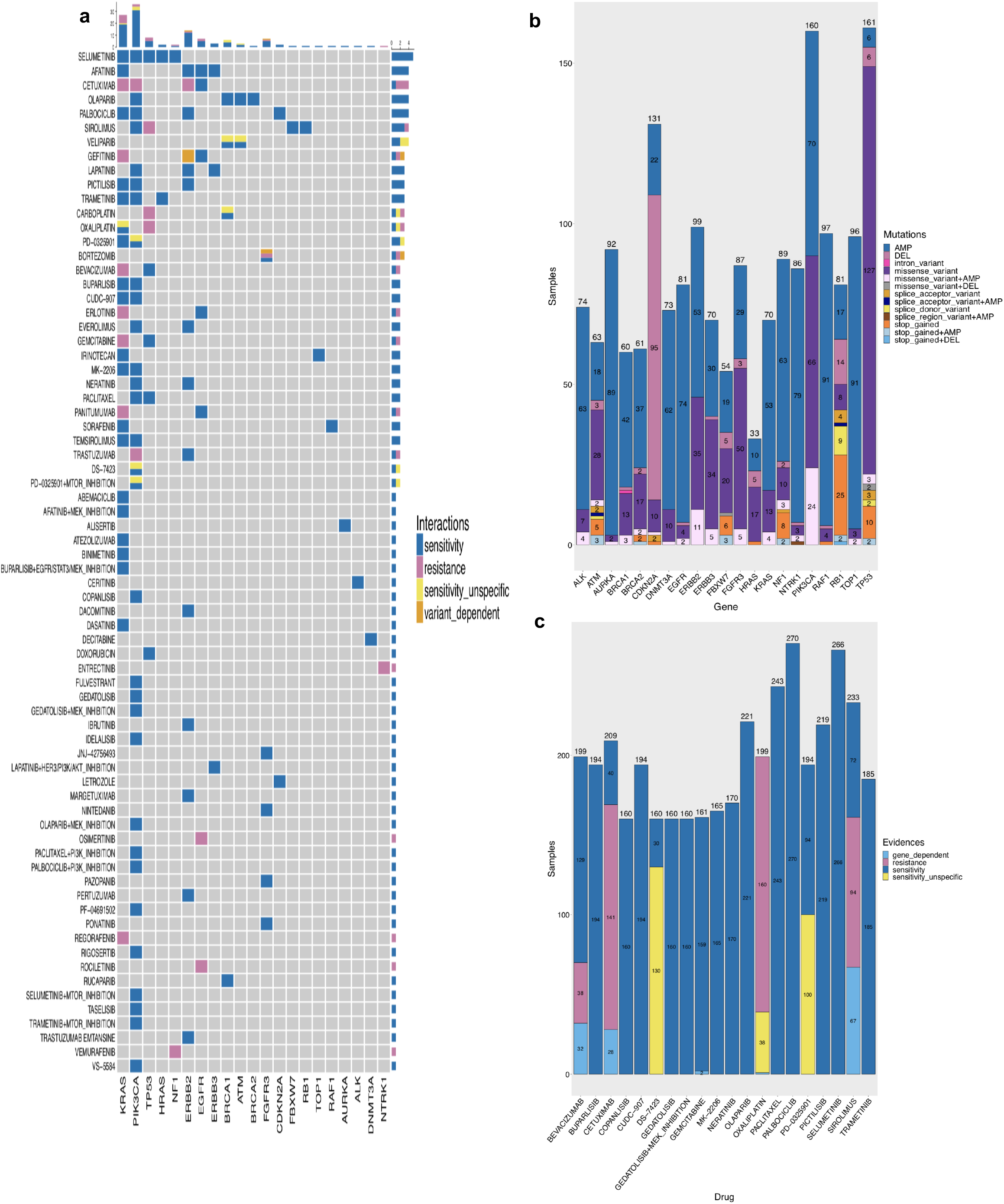
Our drug-prediction workflow discovers a panel of genes and related drugs / drug-combinations for personalized therapy in bladder cancer. **a** The heatmap illustrates the final selection of drug-gene interactions (including combination treatments). “Sensitivity” and “resistance” refer to predicted sensitivity and resistance when the associated gene contains a respective variant. “Sensitivity_unspecific” refers to genes where sensitivity to a particular drug could not be assigned to a particular variant but multiple variants would lead to the prediction. In “variant_dependent” predictions, different variants within one gene can cause a divergent direction of the response prediction (sensitivity/resistance). **b** The barplot shows the prevalence of identified genes and the predicted impact on gene product (abbr.: DEL = deletion, 0 copies; AMP = amplification, > 4 copies). **c** The barplot depicts the 20 most frequently predicted drugs and the directions of drug response. The legend is the same as in **Fig. 5a**, with the exception of “gene_dependent”, which categorizes samples that have multiple genes mutated that have a divergent (sensitivity/resistance) response to the same drug.

The finally selected set represents a mix of genes and drugs known in the context of bladder cancer, together with interesting potential new targets. 45 of the identified 72 drugs or drug combinations are currently or formerly tested in BLCA clinical trials. While being tested in solid tumor basket trials or other solid tumor entities, at the time of our investigation 17 of the listed drugs have not been tested in explicit BLCA trials yet (ceritinib; CUDC-907 (fimepinostat); dacomitinib; DS-7423; entrectinib; gedatolisib; idelalisib; letrozole; margetuximab; MK-2206; PD-0325901 (mirdametinib); PF-04691502; pictilisib; ponatinib; rigosertib; rociletinib; VS-5584).^9^ Twelve predicted drugs are being investigated in the MATCH trial,^10^ five in the My Pathway trial ^11^ and 12 in the COXEN trial.^12^ As expected, only a minority of 12,5% (9/72) of drug approaches are non-targeted therapies.

For ten DGIs reasonable combinational therapies could be identified to avoid therapy evasion. Furthermore, at the time of investigation we found six of the identified combinational therapy approaches in clinical trials with mostly solid tumors (buparlisib + MEK-inhibition; gedatolisib + MEK-inhibition; lapatinib + AKT-inhibition; paclitaxel + PI3K-inhibition; palbociclib + PI3K/mTOR-inhibition; selumetinib + mTOR-inhibition; trametinib + mTOR-inhibition).^9^

The majority of the identified genes (16/21) are currently a subject of clinical trials as diagnostic biomarkers or in reference to therapeutic decision making for BLCA therapy. However, at the time of our investigation DNMT3A, FBXW7, HRAS, RAF1 and TOP1 are not yet found as biomarkers in BLCA related clinical trials and we are the first to report them as potential candidates for biomarker driven therapy testing.^9^ With FGFR3 we identified the first and as yet only genetic biomarker with FDA approval for locally advanced and metastatic BLCA ^13^ amongst our predictions.

### Divergence in drug response prediction

Within a majority of samples, drug response predictions depended on multiple different mutations. In each individual sample a directed response prediction (sensitivity/response) could depend on different genes and therefore divergence of the direction of prediction was possible (in the following referred to as “gene-dependent divergence”). In total, 252 pairs of drug and sample contained any kind of divergence, referring to 158 individual samples with multiple divergent predictions. We note that in a majority of samples 55,3% (228/412) the drug response predictions were free of any divergence. The 252 gene-dependent divergences found across the samples can be categorized into 122 within CIViC predictions, 97 within manual curation, and 33 resulting from differences between CIViC and manual curation predictions (2 in CIViC: resistance / manual curation: support; 31 in CIViC: support / manual curation: resistance), respectively. Thus, in all of these cases multiple genes had been mutated that had different impacts on the drug response. We identified divergent predictions exclusively resulting in reference to 8 drugs (docetaxel; sirolimus; cetuximab; bevacizumab; lapatinib; oxaliplatin; gemcitabine; trastuzumab). Between-source divergences corresponded to 3 drugs (cetuximab; sirolimus; gemcitabine). In total, predictions in only 1,2% (5/412) of samples included between-source divergence (arising exclusively from the curation of different DGIs across the curation sources).

### Potential application of the drug and gene panel

The drug and gene panel resulting from our workflow is a groundwork for a personalized therapy testing approach for BLCA. Investigating the predicted drugs, we observe agents that have been clinically tested in BLCA, such as EGFR-inhibitors like cetuximab. Furthermore, mTOR-inhibitors like sirolimus, agents affecting the RAF/MEK/ERK pathway such as selumetinib, or PARP-inhibitors such as olaparib appear in our panel. With 10 identified drug candidates, PI3K-inhibitors form a major group of which only three (buparlisib; copanlisib; taselisib) are currently in clinical testing explicitly in BLCA.

**Fig. 7** illustrates a blueprint for the future application of our workflow, where we intend to analyze sequencing results from patient derived tumor material. The gene and drug panel can serve as a cost-efficient yet comprehensive molecular diagnostics that is tailored to BLCA tumors. To illustrate an example, in the case of a significant PI3K-mutation (found in 38,8% (160/412) of all samples; see **Fig. 6b**) our algorithm predicted variant-dependent sensitivity to 22 drugs and 6 drug combinations. Additionally, three unspecific gene-dependent predictions for sensitivities were found alongside two predicted resistances. Both resistances refer to EGFR-inhibitors; cetuximab and trastuzumab. Moreover, predictions from our algorithm can be weighted in reference to the level of supporting evidence (only genomic / genomic + DRS information available) (see Fig. 6d). Subsequently, testing of the predicted drug response in patient derived tumor models can confer several implications for future clinical administration (see Fig. 6e). The efficacy of predicted sensitivities from drugs of the same group (e.g. the PI3K-inhibitors buparlisib and gedatolisib) could be assessed preclinically. Similarly, the observed resistance predictions can be utilized as negative controls in testing the aforementioned tumor model. Both in-vitro and in-vivo results would mandatorily be reported to the database, the algorithm was interrogating at the outset, to increase or decrease the power of the underlying DGIs.

**Fig. 7.**
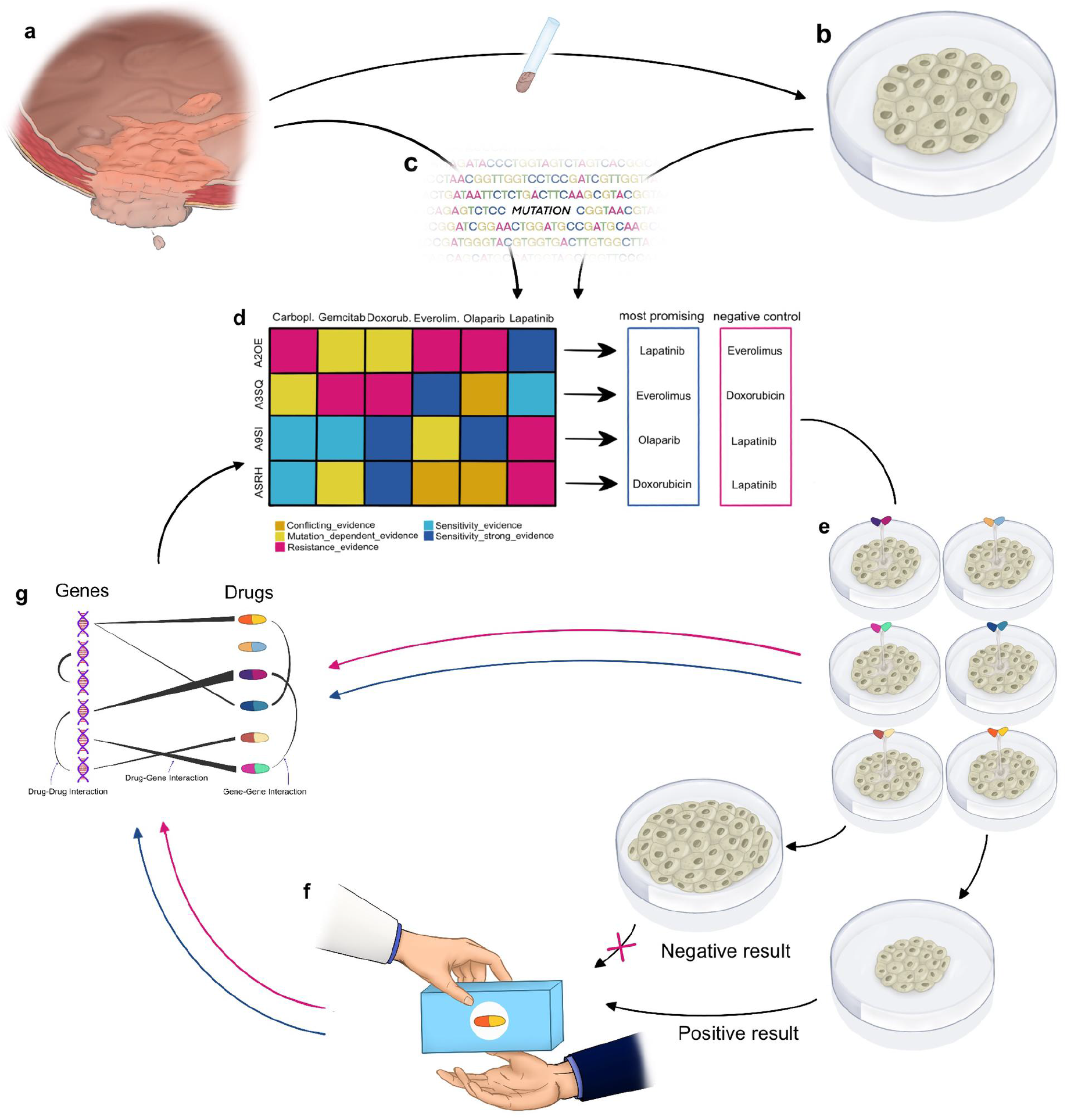
Blueprint for future clinical applications. **a** Histologic samples are derived from the tumor mass and processed into **b** individual patient derived tumor models. **c** Tumor mass and tumor models undergo sequencing to confirm genomic comparability. **d** Sequencing results are analyzed with our algorithm for drug response predictions. Delivering weighted (level of support) and directed (sensitivity/resistance) predictions, the most promising sensitivities can be selected to be tested against predicted resistances, functioning as negative controls. (Figure legend according to **Fig. 5**) **e** The derived proposals are tested in the patient derived models. **f** The drug with the best response is administered to the patient, while feedback about *in-vitro* sensitivities and resistances is reported to the database. **g** Clinical response is also reported to databases (DGIdb; CIViC etc.), increasing or decreasing the power of the DGIs for the next query of the algorithm.

## Discussion

Poor and unforeseeable outcomes of standard therapies in BLCA raise the question, whether empiric “one size fits them all” therapy and testing approaches can be justified in the setting of notably individual tumors.^6^ Indeed, the genomic and transcriptomic diversity in BLCA offers a plethora of known actionable targets.^14^ For other cancer types genomic aberrations have proven to be relevant therapy targets, e.g. BRAF mutations in melanoma or HER2 mutations in breast cancer.^15,16^ In contrast to single marker driven diagnostics and decisions, molecular tumor boards in the US (Mayo Clinic ^17^; Weill Cornell ^18^) as well as in Europe (DKFZ/NCT ^19^; ETH Zürich/SwissMTB ^20^) started to comprehensively investigate individual genomic tumor landscapes. They transferred clinical knowledge amongst oncologic entities resulting in (off-label) drug suggestions for these often end-of-treatment-line patients, with notable responses.^20^ In BLCA, genomic biomarker driven FGFR-inhibitors have only recently been approved.^13^ Despite known targetable mutations, further genomic marker driven therapy has yet to be approved in BLCA. Therefore, BLCA investigations in comprehensive multiple marker based drug prediction are highly warranted.

This study introduces a highly automated algorithm to facilitate a personalized drug response prediction for target-driven therapy testing. To acknowledge the genomic and transcriptomic individuality of BLCA we have developed an algorithm for *in-silico* drug response prediction that includes the highest level of automation reported to date. Our workflow identified 21 genes, corresponding to 72 drugs and drug combinations as candidates for biomarker driven therapy in BLCA. The majority of the identified drugs (45/72) are currently the subject of research for targeted therapy in BLCA and other cancer types. Importantly, even without any *a priori* focus our algorithm identified the single approved genomic marker for targeted therapy in bladder cancer, FGFR3.^13^ This may reinforce that our workflow is a valid approach for the identification of potentially relevant DGIs. Since the method was similarly applied to all genes, we can highlight the discovery of DNMT3A, FBXW7, HRAS, RAF1 and TOP1 as potential candidates for mutation dependent pharmacotherapy, not having been reported in clinical settings up to this point.^9^

Alongside the identification of new potential targets, our WES-based approach could strongly increase the sample coverage. Prior analyses using panel-based sequencing methods identified up to 60% predefined targetable lesions in BLCA patients.^21^ Our WES-based approach identified one or more targetable genetic alterations in 94,7% of muscle-invasive BLCA in the TCGA cohort. Therefore, our algorithm offers a robust strategy for the prediction of promising targets in most patients with muscle-invasive BLCA.

Within the group of identified targetable alterations, we found examples of altered genes playing notable parts in the pathogenesis of BLCA. DNA methylation regulation is crucially dependent on the DNMT3A gene product. Its mutations are described in many cancer entities.^22^ In BLCA, mutated DNMT3A is known to cause hypermethylation and thereby silence promoters of tumor suppressor genes.^23^ So far it is not reported as a biomarker for a targeted therapy approach in BLCA. In the case of DNMT3A activating mutations, our algorithm acknowledged this pathogenetic mechanism and suggested use of the DNA methyltransferase inhibitor Decitabine. The potential importance of combinational therapies in personalized drug discovery became obvious during manual curation. In case of a PIK3CA mutation, our algorithm suggested CDK4/CDK6-inhibitor palbociclib. CDK4/CDK6 is affected downstream in the line of the PI3K (=PIK3CA) pathway, mediated by AKT and Cyclin-D. Due to according references with synergistic efficacy of a combined inhibition of PI3K and CDK4/CDK6 e.g. in breast cancer models, manual curation suggested PI3K-inhibition in combination with palbociclib in case of PIK3CA mutation.^22^

Drug filtering and manual curation after the identification of drug-gene interactions using DGIdb.org uncovered several key findings. We have found that a majority of DGIs were dismissed due to both drug and gene related reasons. Drugs were dismissed due to an absence of evidence as cancer therapies; they were for example investigated in drug metabolism instead. Specialized databases for disease-drug-gene relations would address this problem. Filtering thresholds caused some DGIs to be dismissed due to low cohort prevalence to create a manageable size of a drug and gene panel. However, these dropped DGIs can be of future interest. Ongoing development and increasing the evidence base of expert curated databases such as CIViC will overcome cohort prevalence barriers. During manual curation a majority of non-curated DGIs had to be dismissed due to missing evidence supporting the provided DGI. For instance, mentioning of the drug or gene only in the list of references of a quoted article led to a non-curated DGI. Such DGIs were dismissed because evidence of a single agent gene-dependent drug efficacy was not given.

Automation and introduction of artificial intelligence may solve the problem of vague DGIs. In 2019 Kim et al. introduced the artificial intelligence based search engine for disease-gene-chemical relationships DigChem, which is able to avoid aforementioned text-mining errors. However, their study emphasized that the overlap with other DGI databases (CTD ^23^, IBM Watson for Drug Discovery ^24^ and DrugBank ^25^) is complementary, rather than displaying directly comparable results.^26^ Moreover, in the process of DGI filtering we found a large disparity between provided DGIs and the ratio of expert curated DGIs. Therefore, we conclude that manual curation has still to be considered mandatory to achieve evidence-based DGIs. Although only in a minority of samples, we identified divergent drug response predictions due to the presence of multiple DGIs per sample and due non-matching genome- and transcriptome-based predictions. A majority of divergences resulted from drugs with multiple-gene dependent response predictions within a given sample. In these cases model and clinical testing will facilitate judgment about the dominant DGI. Importantly, at the time of investigation a majority of samples were free of divergences in drug response prediction.

Both curations, manual and expert based, identified more sensitivities than resistances in DGIs. Although validation is missing so far, we credit this fact to a known reporting bias for positive study results eg. sensitivities.^28^ For the creation of comprehensive DGI databases a more liberal reporting of resistances will be essential. Siu et al. have addressed the need for liberal and barrier-free reporting of DGIs in creating a scheme for responsible sharing of cancer genome data for utilization in comprehensive analysis.^29^ Therefore, testing and reporting of drug resistance in DGIs as comprehensive as existing reporting of drug sensitivities, will be of paramount importance.

Biologically, advanced tumor stages are characterized by the consequences of multiple prior therapies in the form of progressed intratumor heterogeneity. Taking tumor evolution theories into account a plethora of potential therapy evasion mechanisms might have developed by that time and create limitations to personalized therapy so far.^30^ However, our algorithm can be adapted for testing in patient derived tumor models and thereby offers a tool to facilitate testing in multiple tumor stages and of multiple locations (eg. metastases). The remarkable findings of Biswas et al. (Nature Medicine, 2019), demonstrated mutational similarities between different evolutionary tumor branches. These findings might open the door to solutions how to address intratumor heterogeneity with targeted therapies.^31^ In our opinion, developing a dynamic understanding of oncogenesis and its underlying processes of therapy evasion development is compelling. It will allow for multi-time and multi-location evaluation of tumors and lead to accordingly individualized therapies.

We acknowledge that our approach is still hypothesis-generating and further validation is needed. Eventually, clinical trials will need to prove the superiority of *in-silico* drug response predictions. In 2019 preliminary data from the BISCAY trial compared the efficacy of the PD-L1 checkpoint inhibitor durvalumab in combination with targeted therapies in selected urothelial cancer patients. Despite achieving a superior overall response rate with the combination of durvalumab and the PARP inhibitor olaparib in case of an DNA-damage-repair gene alteration, compared to durvalumab monotherapy none of the study arms could reach the threshold for a positive study result.^32^ In 2017 Kiss et al. demonstrated that genomic variations and their impact as predictive markers for therapy response, have to be carefully interpreted with consideration of transcriptional and gene-regulatory factors as well as of co-existing mutations.^33^ These findings highlight the importance of biomarker-driven trials. Basket-trials such as the My Pathways study (<u>NCT02091141</u>) or the NCI-MATCH study (<u>NCT02465060</u>) offer access to individualized treatment with a diversity of targeted agents.

Our algorithm represents a sophisticated tool that can be implemented in clinical trials and lay the foundation to precise cancer therapy based on a comprehensive understanding of molecular oncogenesis.

In conclusion, our algorithm for *in-silico* drug response prediction in bladder cancer identified drugs with proven antitumor activity but importantly also new potentially druggable targets and novel drugs, which have not yet been tested in bladder cancer. Moreover, our drug prediction provided information for 94,7% of BLCA samples, highlighting its applicability to the majority of bladder cancer patients. Integration of genomic and transcriptomic data acknowledged different regulatory processes in oncogenesis and increased the granularity of our predictions. Importantly, our algorithm yielded an efficient and affordable gene panel for drug response prediction based on a comprehensive approach. Integration of proven and disproven drug-gene interactions into open source databases will be of utmost importance to improve drug predictions. Prospectively, our workflow will include these growing databases and consequently predict promising drugs contemporaneously. Ultimately, these *in-silico* drug response predictions require further testing in patient derived models and clinical trials prior to implementation as personalized approaches in clinical practice.

## Methods

### The Cancer Genome Atlas bladder cancer cohort

We analyzed the genomic landscape of the 412 bladder cancer patients of the TCGA-BLCA cohort ^14^. Based on the whole exome information, variants in the tumor genome have been identified and linked to their potentially targeting drugs. We prioritized variants and drugs according to their clinical relevance and prevalence in bladder cancer patients. The workflow is depicted in Figure 1 and detailed in the following. The 412 patients in the TCGA-BLCA cohort present a well-mixed population, e.g. in regard to gender, BLCA subtype, and tumor stage, which will likely reduce the effect of clinical features on the drug prediction results.

For each patient in the TCGA-BLCA cohort, the whole exome sequencing based bam files for normal as well as tumor samples have been downloaded from the GDC data portal.^34^ Bam files had already been generated with the GDC data harmonization workflow.^35^ For patients with multiple tumor and/or normal samples, we selected the sample with the latter plate number according to the GDC guidelines ^36^, and preferred fresh frozen over FFPE samples. Further, to ensure a sample selection as homogeneous as possible, we selected the blood derived normal whenever possible, as this was the normal control available for the majority of patients.

### Variant Calling from the TCGA cohort

To investigate the genomic landscape of the 412 bladder cancer patients, we performed somatic variant calling for the identification of single nucleotide variants (SNVs), as well as copy number calling to identify copy number variants (CNVs). The pipeline for whole exome sequencing analysis from the retrieved bam files to unannotated variant calls is based on the framework described in J. Singer et al. 2018 ^37^, employing the Snakemake workflow environment ^38^

#### SNVs

In brief, variants were called based on the GATK best practices workflow ^39^, with somatic variant calling on the matched tumor and normal samples. To achieve high quality calls and reduce the number of false positives, we applied three different variant callers and only considered variants that were identified by at least two callers. The utilized variant callers were MuTect ^40^, Strelka ^41^, and VarScan2 ^42^, where the latter two not only perform single nucleotide variant calling, but also identify small insertions and deletions in the sample. Variants were annotated using SNPeff ^43^ and SNPsift ^44^ and enriched with information on their presence in ClinVar ^45^, COSMIC ^46^, and dbSNP ^47^, as well as their overall mutation impact (e.g. missense variant or synonymous variant). We also performed a functional annotation to assess the potential of each variant to be damaging for the protein function.

#### CNVs

We used the CNV caller Facets ^48^ to call copy number variants on the matched tumor and normal samples for each patient. All CNVs were annotated to identify the affected genes.

### *In-silico* drug prediction and selection

Similarly as described in Singer et al. ^20^, we queried DGIdb ^7,49,50^ to identify the drug-gene interactions reported in a collection of 30 databases, of which several are expert-curated (e.g. MyCancerGenome^51^). The more databases support an interaction, the higher the scoring of the drug. Finally, based on drug-gene interactions identified in DGIdb we collected associated clinical trials at ClinicalTrials.gov. Here we distinguished between clinical trials in BLCA and those also including solid tumors (cancer type, “ct”), non-cancer type specific (“nct”) trials, and trials not related to cancer in general (“other”). Trials in the “nct” category focus on patients with other cancer types, thus for bladder cancer patients their corresponding drugs would potentially be regarded as off-label therapy.

Integrating clinical trial information provides a first prioritization of the drug-gene interactions (DGIs) resulting from the DGIdb query. The initial set of possible drugs was filtered to only contain drugs that have previously been tested in cancer-related clinical trials (categories “ct” and “nct”). Further, we prioritized drugs according to their frequency in the cohort, to obtain a set of drugs that is likely to contain at least one possible match for each BLCA patient.

### Variant prioritization and selection

The set of identified variants (both SNVs and CNVs) was filtered to prioritize variants that are likely to be of clinical relevance, i.e. that i) are likely to have an impact on the protein function and ii) that are associated with genes with DGIs listed in DGIdb. An overview of the filtering process is illustrated in Figure 1. SNVs were filtered to only include those that have a high impact (higher than 20, based on the SNPeff impact list) and that are predicted to be damaging. The CNVs were filtered to only include those with deep deletions (Copy number = 0) or high amplifications (Copy number > 4).

Further, we only included variants that affect genes with a reported drug-gene interaction (based on the DGIdb query). Furthermore, we prioritized variants that occur more frequently across the TCGA-BLCA cohort, to obtain a final selection that is not only likely to be of clinical relevance, but also likely to have at least one targetable variant present in BLCA patients.

The minimum set cover analysis was performed to identify the fraction of the cohort that was covered by the selected set of priority genes (and drugs). It was based on in-house scripts and performed a greedy selection of the best possible set.

### Integration of CIVIC

The information provided by DGIdb is very comprehensive and thus a valuable resource to provide a basic set of DGIs as a starting point. However, it has several limitations that impede its direct use for clinically relevant *in-silico* drug prediction. First of all, it is undirected and only lists DGIs without indicating the nature of the DGI, i.e. whether a mutation in the respective gene confers sensitivity or resistance. Second, it only reports interactions between genes and drugs and does not consider the individual variants present in a patient’s tumor. And finally, it contains not only expert-curated databases such as MyCancerGenome.org, but also comprehensive compound collections. As a consequence, we enriched the DGIdb based information with variant specific expert curated information from the CIViC database.^8^ Here, individual aberrations that affect a specific gene are evaluated for their potential to confer sensitivity or resistance to a drug, and each evaluation (called evidence item) is rated with an evidence level to show the quality of the underlying information. For instance, preclinical studies are rated lower than information gained from multi-center clinical trials. Our CIViC based assessment of each DGI is accordingly applied on the variant level, such that in case different TCGA samples showed different variants of the same gene, a DGI can have divergent readouts, i.e. sensitivity or resistance. Note that it is possible that a variant-drug interaction type is not clearly identifiable from the information currently available in CIViC, e.g. in some studies a variant seemed to confer sensitivity while in other studies this was not a significant finding. In these “CONFLICT” cases we employ a majority vote that counts how many evidence items support sensitivity and resistance, respectively, and take the interaction type supported by the majority of evidence items. Note that before the majority vote is applied, we prioritize the evidence items according to their relevance, e.g. evidence items on bladder cancer would be prioritized over those with information from other cancer types. Further, CIViC contains evidence items that are not explicitly in favor of either sensitivity or resistance. Accordingly, the corresponding DGI is labeled as “UNKNOWN” to indicate that the DGI was found in CIViC, but that the available information is not sufficient to clearly predict response or resistance.

### Manual curation of drug-gene interaction

References of DGIs obtained from the DGIdb query with no annotation in the curated CIViC database were selected for manual curation. Clinical studies as well as experimental studies were included. First, categories for response were separated into “SENSITIVITY”, “RESISTANCE” and “UNKNOWN”. The category “UNKNOWN” thereby summarized all references with missing evidence of single agent activity, effectiveness or missing reference to the drug or gene itself. An annotation category was added to document restrictions such as: quality of data, conflicting evidence, variant dependent response, co-variant dependent response, gene expression dependent response, reasonable combinational therapy to avoid therapy evasion and clinically relevant information about the drug. Second, the full text articles referenced by DGIdb were screened for the gene and the active substance. If the active substance name was not found, generic names, brand names and accession numbers were searched for.

If gene and drug were found, data was screened to support single agent activity of the drug in clear reference to an explicit mutation and/or wild type. In clinical studies response “SENSITIVITY” was defined as improvement of response rate, progression of free survival or overall survival under administration of the drug, in case of mutated gene compared to wild type. Response “RESISTANCE” was here defined as decreased response rate, inferior progression free survival or overall survival under administration of the drug, in case of a mutated gene compared to wild type. In experimental studies, response “SENSITIVITY” was defined as deteriorated growth or decreased survival of the tumor model under administration of the drug, in case of mutated gene compared to wild type. Response “RESISTANCE” was defined as accelerated growth or increased survival of the tumor model under administration of the drug, in case of mutated gene compared to wild type. Response “UNKNOWN” was defined as no significant difference of clinical outcome or tumor model reaction when comparing mutant to wild type. References with missing evidence of single agent activity, effectiveness of the administered drug or missing reference to the drug or gene itself, were classified as response “UNKNOWN” as well.

### Integration of DRS

We used the CellMiner tool to mine for drug response correlating to expression data of the NCI-60 cell lines.^52^ As previously published, corresponding genes were used to generate patient specific drug response scores (DRS) using correlation coefficients as weighting factors.^53^ Thereby, DRS represent a statement on a potential drug response based on the level of expression of a certain gene product. Hence, a positive DRS is a prediction of a drug sensitivity, as the underlying investigations correlated tumor model response to a certain level of gene product expression. However, a negative DRS does not predict a therapy resistance, but a non-response to the drug. A therapy “RESISTANCE” is defined in **Methods - Manual curation of drug-gene interaction**. An overview of the DRS predictions for the TCGA samples is shown in **Fig. 5**.

## Data availability

The data that support the findings of this study are available from the corresponding author upon request.

## Author contributions

R.S., F.S., M.L.R.G. and F.K. designed and supervised the development of the study workflow. F.S. and M.L.R.G. performed variant calling and annotation of variants. F.K., F.S., M.L.R.G. and R.S. performed analysis and interpretation. F.K. performed manual curation. F.K., F.S., M.L.R.G. and R.S. drafted the manuscript. F.K. and N.K. designed and N.K. illustrated Fig. 6. F.K., F.S., M.L.R.G., E.A.G., Y.L., E.D., M.K.J. and R.S. performed manuscript editing.

